# Perilacunar bone tissue exhibits sub-micrometer modulus gradation which depends on the recency of osteocyte bone formation

**DOI:** 10.1101/2021.09.21.461298

**Authors:** Caleb J. Rux, Ghazal Vahidi, Amir Darabi, Lewis M. Cox, Chelsea M. Heveran

**Affiliations:** Department of Mechanical & Industrial Engineering, Montana State University; UC Berkeley-UCSF Graduate Program in Bioengineering

**Keywords:** Atomic force microscopy, osteocyte, lacunae, lacunar-canalicular remodeling, aging, bone

## Abstract

Osteocytes are capable of resorbing and replacing bone local to the lacunar-canalicular system (LCS remodeling). However, the impacts of these processes on perilacunar bone quality are not understood. It is well established that aging is associated with reduced whole-bone fracture resistance, reduced osteocyte viability, and truncated LCS geometries, but it remains unclear if aging changes perilacunar bone quality. In this study, we employed atomic force microscopy (AFM) to quantify sub-micrometer gradations from 2D maps surrounding osteocyte lacunae in young (5 mo) and aged (22 mo) female mice. AFM-mapped lacunae were also imaged with confocal laser scanning microscopy to determine which osteocytes had recently deposited bone as determined by the presence of fluorochrome labels. These assays allowed us to quantify gradations in nanoscale mechanical properties of bone-forming/non-bone-forming osteocytes in young and aged mice. This study reports for the first time that there are sub-micrometer gradations in modulus surrounding lacunae and that these gradations are dependent upon recent osteocyte bone formation. Perilacunar bone adjacent to bone-forming osteocytes demonstrated lower peak and bulk modulus values when compared to bone near non-bone-forming osteocytes from the same mouse. Bone-forming osteocytes also showed increased perilacunar modulus variability. Age reduced lacunar size but did not significant effect modulus gradation or variability. In general, lacunar morphology was not a strong predictor of modulus gradation patterns. These findings support the idea that lacunar-canalicular remodeling activity changes the material properties of surrounding bone tissue on a sub-micrometer scale. Therefore, conditions that affect osteocyte health have the potential to impact bone quality.

**GRAPHICAL ABSTRACT:** 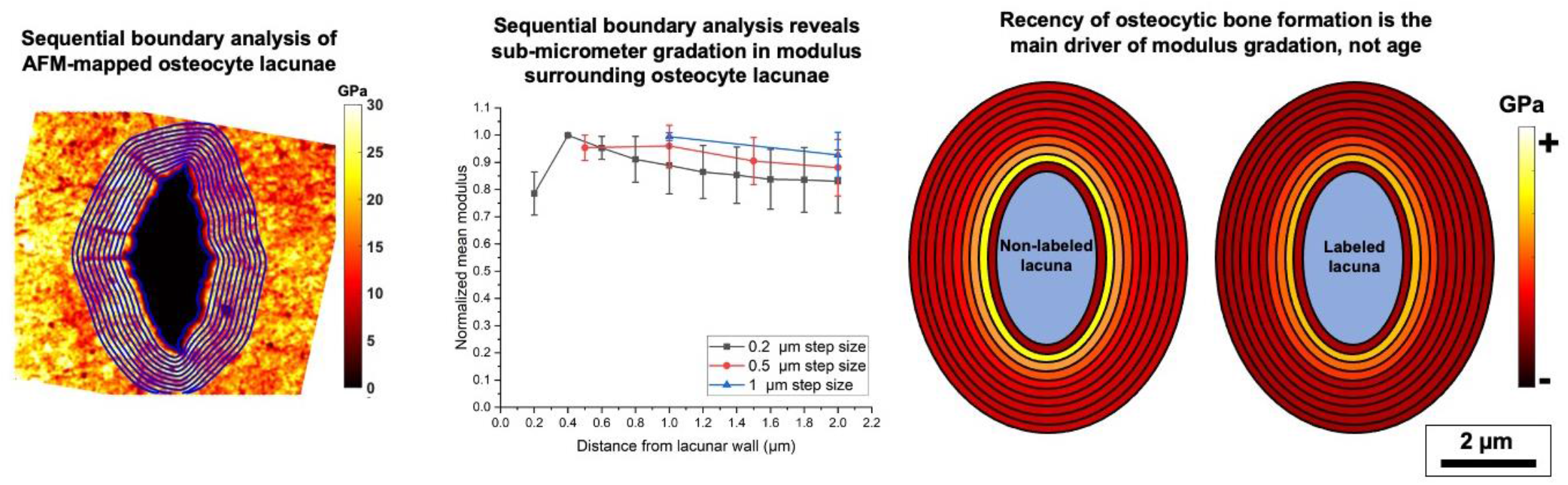

## INTRODUCTION

Osteocytes, the most common cells in bone, live in a dense interconnected network of micrometer-scale voids in mineralized bone tissue (lacunae) connected by sub-micrometer-radius channels (canaliculi) [1–7]. The lacunar-canalicular system (LCS) has an estimated 215 m^2^ surface area in the human skeleton [7] and its trillions of connections allow for osteocytes to communicate within the skeletal network and to other systems in the body [3,5,7,8]. Osteocytes can modulate the size of the LCS through either resorbing bone [9–13] or replacing new osteoid [1,10,14–20] in a process termed LCS remodeling. This process contributes to systemic calcium homeostasis, as demonstrated by expanding lacunae and canaliculi in lactation and recovery after weaning [10,14,21]. It is not yet understood if LCS remodeling also contributes to the maintenance of bone quality. This question is of importance because LCS geometries truncate in aging [2,20,22–29], which suggests that aging alters LCS remodeling activity.

Prior work suggests that LCS remodeling has the potential to reduce bone quality. In DMP1^Cre−/−^ mice and MMP-13^−/−^ mice, both phenotypes demonstrate reduced LCS remodeling activity together with reduced notched fracture resistance [30,31]. However, it is not clear why bone with less LCS remodeling has lower bone fracture resistance. It is possible that morphological changes to the LCS affect the tendencies of cracks to initiate and propagate. On the other hand, it is possible that LCS remodeling directly benefits bone quality. Improvements in bone quality with LCS remodeling could result from a decrease in tissue maturity, leading to bone with lower mineralization and modulus in the vicinity of lacunae and canaliculi [32]. Several previous studies report sub-micrometer-scale mass gradation near lacunae and canaliculi, where the least mineralized region is found within the first few hundreds of nanometers of bone tissue adjacent to lacunar and canalicular walls [1,33]. However, it has not been evaluated whether the gradations in bone material properties near the LCS are influenced by the remodeling activity of the osteocyte.

It is possible to evaluate bone quality near bone-forming and non-bone-forming osteocytes by using fluorochrome labeling and high-resolution material property mapping. Fluorochrome labels are small enough to travel through the LCS and are observed to label lacunae [34–39]. While they may also label canaliculi, most confocal techniques lack the appropriate resolution to discern these smaller features. Bone quality adjacent to lacunae has usually been evaluated with conventional micrometer-scale-resolution characterization tools (i.e., Raman spectroscopy, nanoindentation, quantitative backscattered scanning electron microscopy), but these techniques fail to identify variation near lacunae outside of extreme phenotypes [6,14,32,40–45]. The lack of detection of material property variation near lacunae is a reflection of the resolution of these tools, since line profiles collected through synchrotron-based techniques demonstrate mass gradation near lacunae and canaliculi on the scale of hundreds of nanometers away from LCS walls [1,33]. To date, the spatial variation of bone quality near the LCS has not been generated in 2D maps and has not been compared for bone-forming and non-bone-forming osteocytes.

Atomic force microscopy (AFM) is well-suited for mapping bone quality near bone-forming and non-bone-forming osteocytes. AFM quantitatively assesses modulus on the order of 10s of nanometers using fast force mapping techniques. AFM has been used to demonstrate that modulus is heterogeneous near lacunae and canaliculi in 4-month female Wistar rats, although the specific gradation of modulus with respect to distance from the LCS was not evaluated [6]. Several challenges exist to analyzing modulus maps of lacunae for these spatial data, including reliably defining and smoothing the lacunar edge for a variety of lacunar shapes and sizes, sequentially expanding the lacunar edge by a given step size to create analysis regions (e.g., pixels with set range of distances from the lacunar wall), and determining appropriate step sizes for resolving modulus gradation with distance from the lacunar wall. Thus, we sought to identify an analytic approach to analyzing perilacunar modulus maps for the purpose of comparing bone quality between bone-forming and non-bone-forming osteocytes.

The purposes of this study were to (1) develop an approach to analyze AFM-generated modulus maps of perilacunar bone for 2D spatial gradation, (2) determine whether labeled lacunae have different perilacunar modulus variation than non-labeled lacunae, and (3) evaluate whether aging impacts perilacunar modulus variation. To develop the analytic approach and investigate our research question, we utilized skeletally-mature young adult (5 mo) and early old age (22 mo) female C57Bl/6 mice, since this mouse model and age range produce marked changes in LCS morphology [19]. We hypothesized that lacunae would demonstrate modulus variation in agreement with mineralization variation reported with high-resolution techniques, that labeled lacunae would have lower moduli than non-labeled lacunae, and that aging would decrease the size of the region of lower-modulus bone near lacunae.

## RESULTS

We first developed AFM mapping and analysis techniques in order to determine whether perilacunar modulus demonstrates gradation with respect to distance from the lacunar wall and at what resolution this gradation is apparent. We then used these mapping and analysis parameters to investigate the influence of osteocyte bone formation activity on perilacunar modulus in skeletally mature (5 mo) and early old age (22 mo) female C57Bl/6 mice.

### Perilacunar bone tissue shows submicrometer-scale gradation in modulus

Atomic force microscopy fast force mapping demonstrates that bone modulus has submicrometer-scale gradation adjacent to osteocyte lacunae in cortical bone of the murine femur (**Figure 1a**). To assess the effect of distance from the lacunar wall on modulus, we initially obtained eight maps from one 7-month female C57Bl/6 mouse. For each map, we binned pixels within regions of three different step sizes, 0.2, 0.5, and 1μm, extending outward to 2 μm from the lacunar wall. The smallest step size, 0.2 μm, was selected because this distance is greater than the smallest lacunar spatial features but does not reduce the number of pixels per ring to such a low level as to impede interpretation of histograms. Further, gradations in mass density from synchrotron line profiles occur at a similar length scale [33]. We also studied 0.5 μm and 1 μm step sizes (i.e., averaging over all pixel modulus values within concentric rings of this width), since these resolutions are close to those of other common bone quality measurement techniques (e.g., Raman spectroscopy, backscattered scanning electron microscopy, nanoindentation). At each distance, a mean and a standard deviation were calculated from a histogram of all pixels within the region (**Figure 1**).

**Figure 1.**
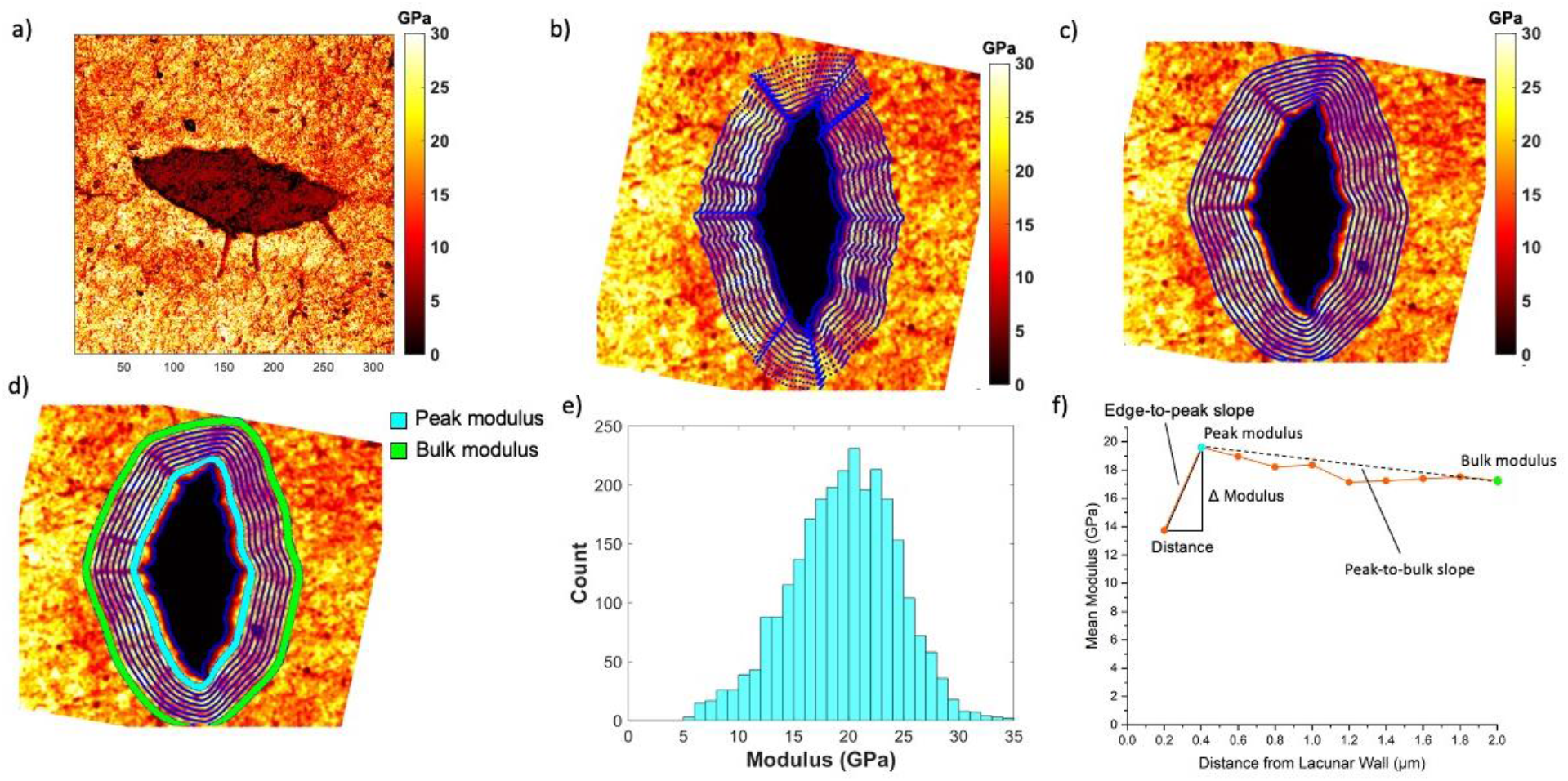
a) An AFM modulus map for an osteocyte perilacunar bone. b) Raw modulus maps are processed through masking, rotation, and dilation steps. Sequential concentric rings are assigned for analysis. In this image, concentric rings are distanced by 0.2 μm. c) A convolution operation smooths boundaries to identify the lacunar wall. d-e) All pixels for an individual concentric ring, such as shown in cyan, are used to construct a histogram (bin size 1 GPa) of moduli. f) The modulus versus distance gradation profile corresponding to mean modulus values found within sequential concentric ring regions (cyan indicating the region that contains the peak mean modulus, green indicating the region that contains the bulk mean modulus value).

Analysis of all maps at each of the three step sizes demonstrates that step size influences the ability to discern modulus gradation (**Figure 2a**). At a step size of 0.2 μm, the modulus rose to a peak at 0.2-0.4 μm from the lacunar wall and then declined towards a bulk bone (i.e., 2 μm from the lacunar wall) (**Figure 3**). These gradations were apparent in both raw data and data normalized to a peak value per lacunar map. The larger step sizes of 0.5 μm and 1μm failed to capture the rise to a peak and decline to bulk seen in mean modulus values when using a finer 0.2 μm step size (**Figure 2a**). Standard deviation was also evaluated at each step size. Using a 0.2 μm step size, standard deviation was found to be greatest close to the lacunar wall and declined within 0.4-0.6 μm to stable values (**Figure 4**). However, standard deviation has less sensitivity to step size. All three step sizes detected a decrease in standard deviation with distance from the lacunar wall, although the resolution of this effect improves with finer step size (**Figure 2b**).

**Figure 2.**
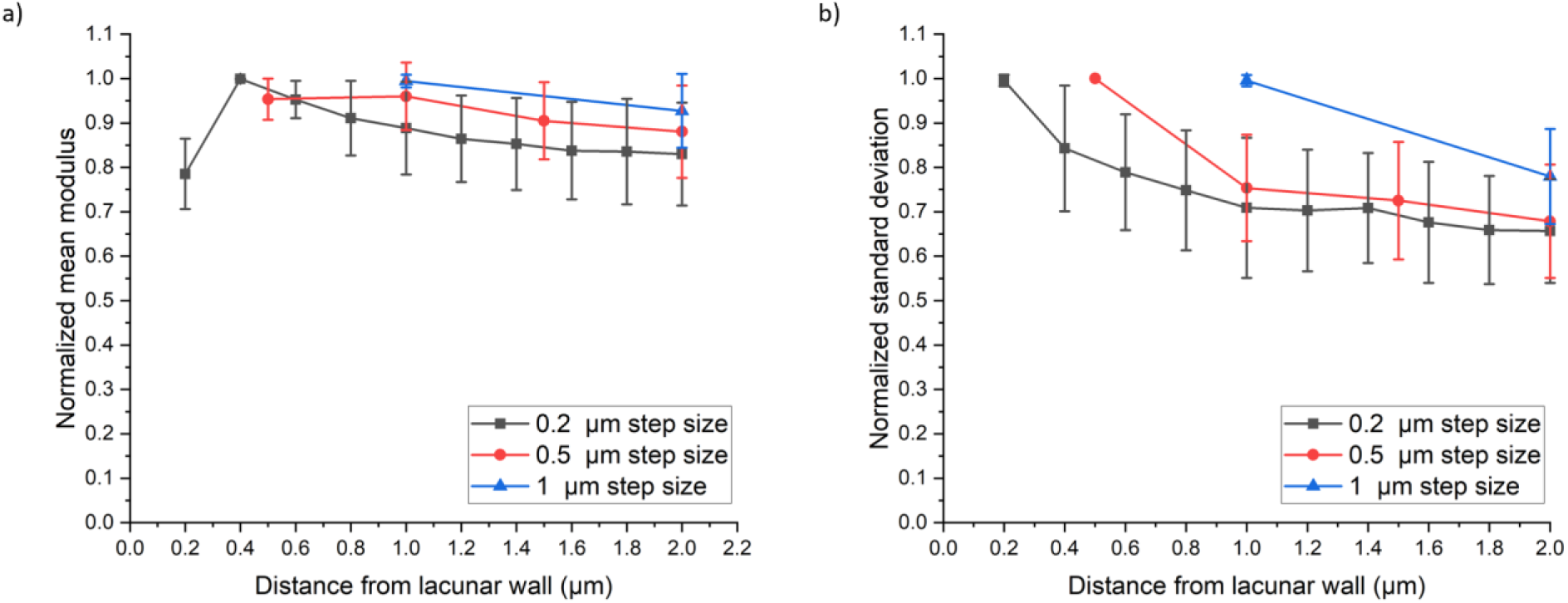
a) Normalized mean moduli versus distance from the lacunar wall is plotted with data from 0.2, 0.5, and 1 μm step sizes extending to 2 μm from the lacunar edge. The distance from the lacunar wall indicates the outer distance of a bin (e.g., 0.4 μm means 0.2 – 0.4 μm). Error bars represent one standard deviation. b) Normalized standard deviations versus distance from the lacunar wall is plotted with data from 0.2, 0.5, and 1 μm step sizes extending to 2 μm from the lacunar edge. Error bars represent one standard deviation. Plots created from eight AFM maps obtained from lacunae from one 7-month female C57Bl/6 mouse.

**Figure 3.**
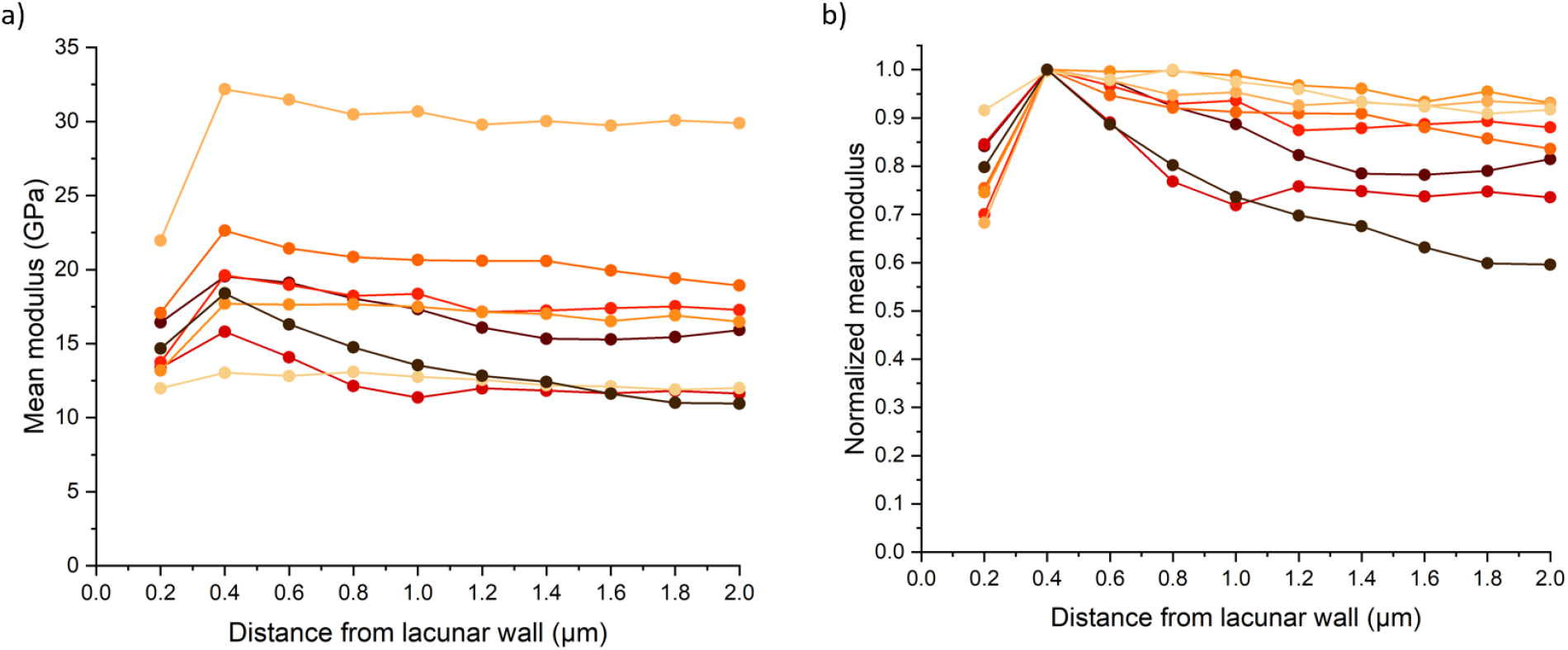
a) Mean modulus for each concentric ring plotted against distance from the lacunar wall. The distance from the lacunar wall indicates the outer distance of a bin (e.g., 0.4 μm means 0.2 – 0.4 μm). Connected dots each represent individual osteocyte lacuna map. b) Normalized mean modulus for each concentric ring plotted against distance from the lacunar wall. Mean modulus values were normalized against the peak mean modulus value for a given map. Plots created from eight AFM maps obtained from lacunae from one 7-month female C57Bl/6 mouse.

**Figure 4.**
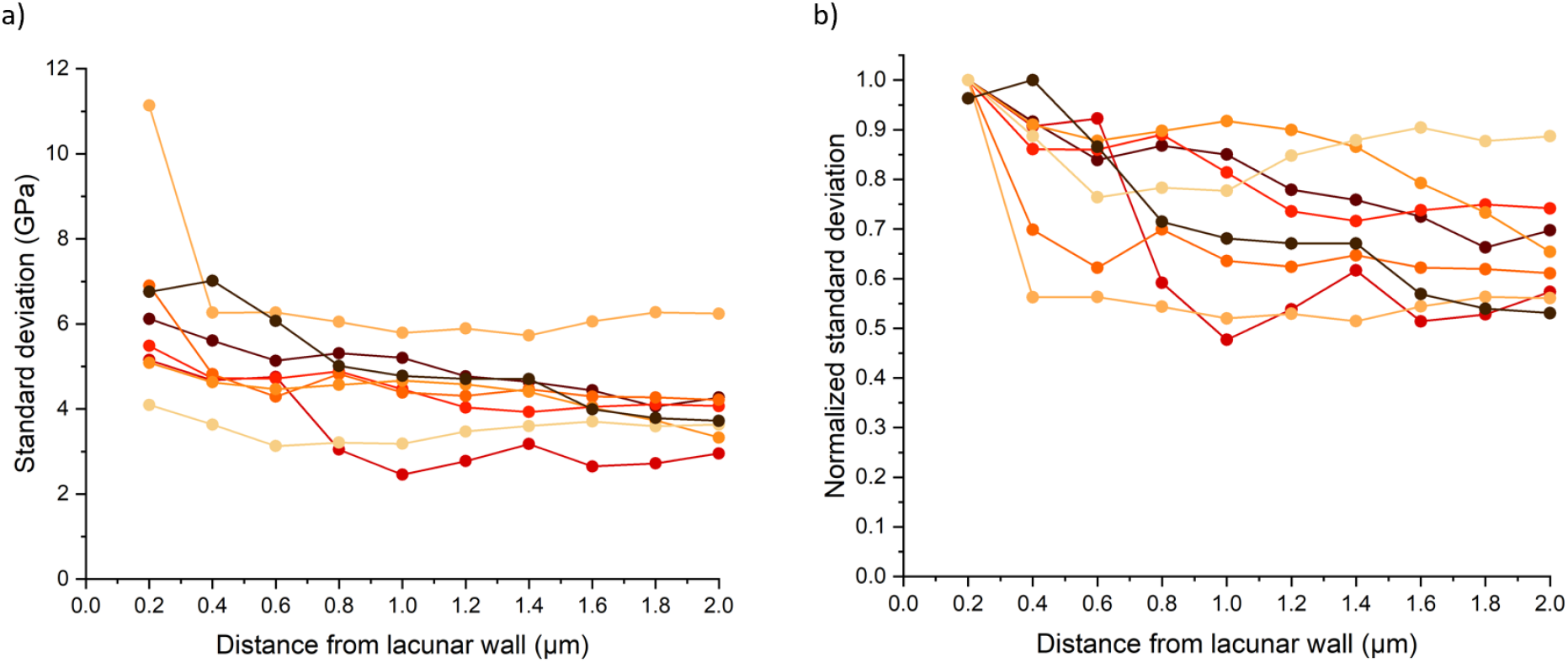
a) Mean standard deviation for each concentric ring plotted against distance from the lacunar wall. The distance from the lacunar wall indicates the outer distance of a bin (e.g., 0.4 μm means 0.2 – 0.4 μm). Connected dots each represent individual osteocyte lacuna map. b) Normalized standard deviations for each concentric ring plotted against distance from the lacunar wall. Standard deviation values were normalized against the peak standard deviation value for a given map. Plots created from eight AFM maps obtained from lacunae from one 7-month female C57Bl/6 mouse.

### Bone-forming osteocytes have distinct perilacunar modulus gradation compared with non-bone-forming osteocytes

To identify whether LCS bone formation and age influence bone tissue modulus, we first sought to identify bone-forming lacunae. Through confocal laser scanning microscopy (CLSM) imaging, we determined that either calcein or alizarin administered 2 days before euthanasia to 5 mo and 22 mo female C57Bl/6 mice abundantly labeled cortical femur osteocyte lacunae (**Figure S1**). Labels administered at 8 days before euthanasia were infrequently found. Together, these findings suggest that osteocytes frequently deposit new osteoid and that this bone tissue undergoes frequent turnover.

We then evaluated whether osteocyte perilacunar remodeling affects bone tissue modulus gradation by comparing modulus gradation between lacunae in the femur anterior quadrant for osteocytes that were forming bone (alizarin-labeled) or not forming bone (no label) (**Figure 5**). Of the five lacunae per mouse randomly selected for AFM mapping, 60% showed an alizarin bone label (administered 2 days before euthanasia) in both 5 mo and 22 mo mice (n = 5 mice for each group). None of the mapped lacunae were labeled with calcein (administered 8 days before euthanasia). Mixed model ANOVA showed that labeled lacunae had lower peak modulus (− 11.72%, p < 0.05) and bulk modulus (−10.06%, p < 0.05) (**Table 1**). There were no interactions between label and age for these measures. Of note, several labeled lacunae had much greater distance to the peak mean modulus. However, on average, the distance to peak mean did not differ between labeled and non-labeled lacunae. Labeled lacunae also had decreased peak standard deviation (−11.06%, p < 0.05) and bulk standard deviation (−12.61%, p < 0.05).

**Table 1:**
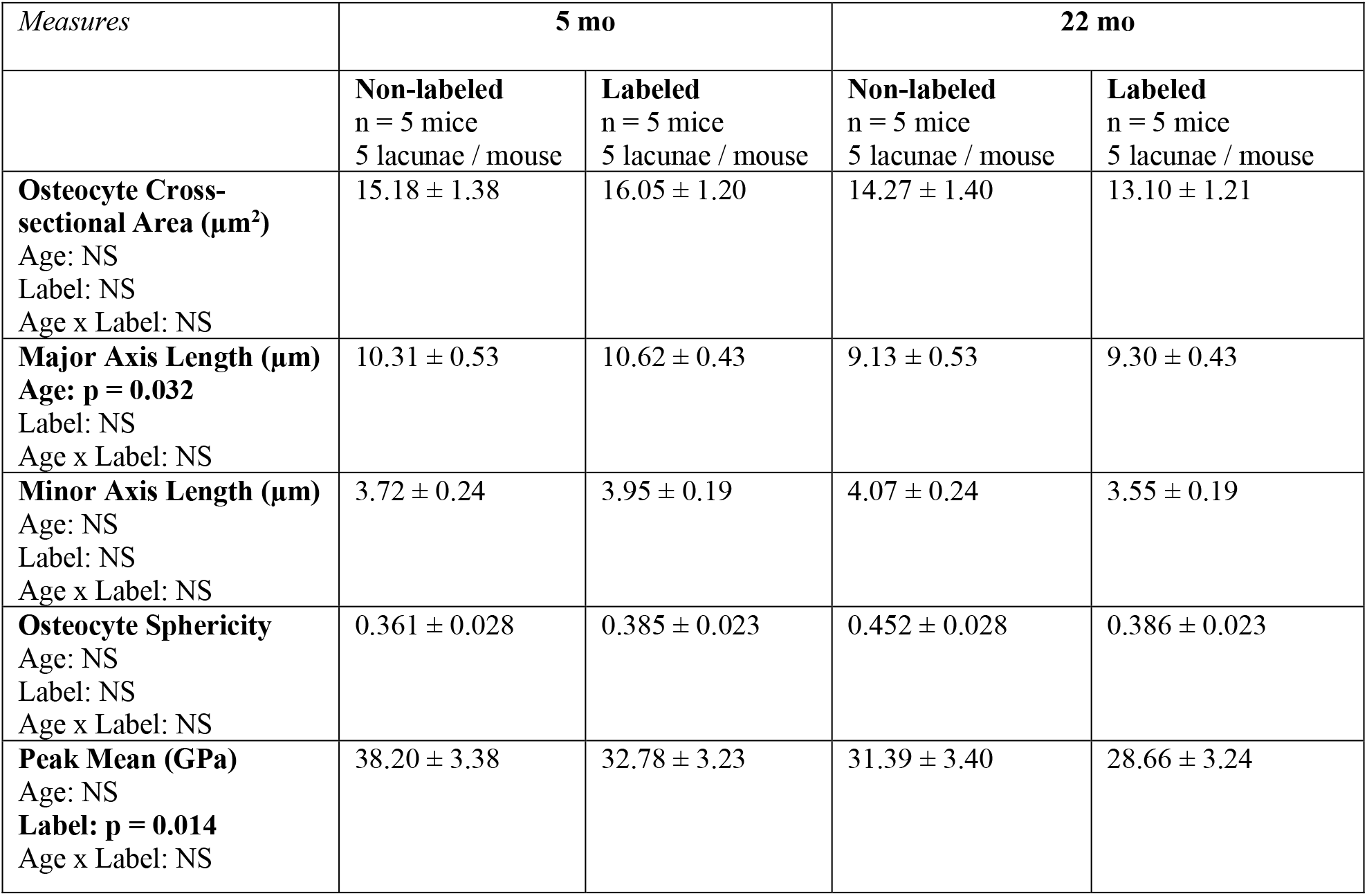
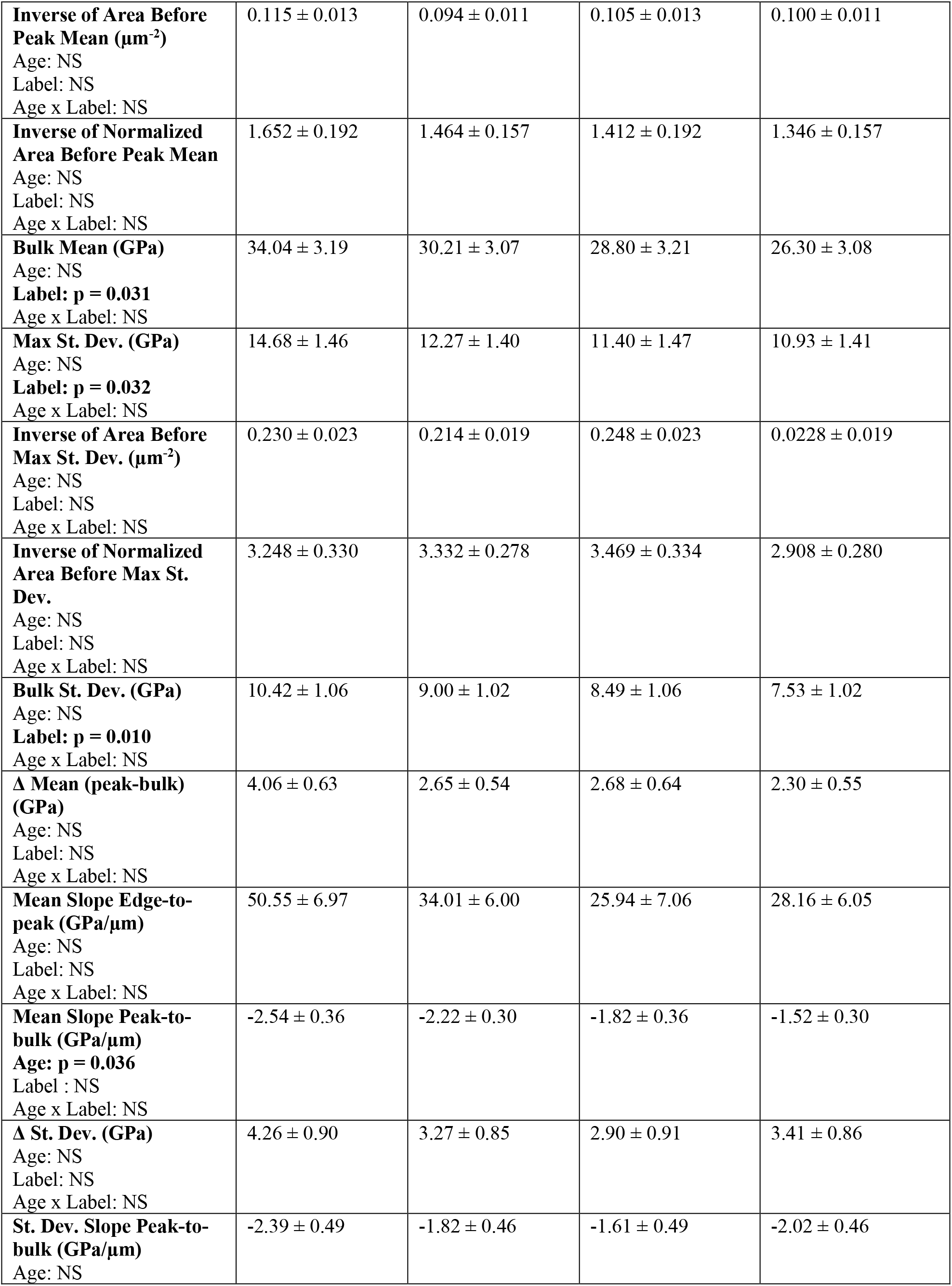
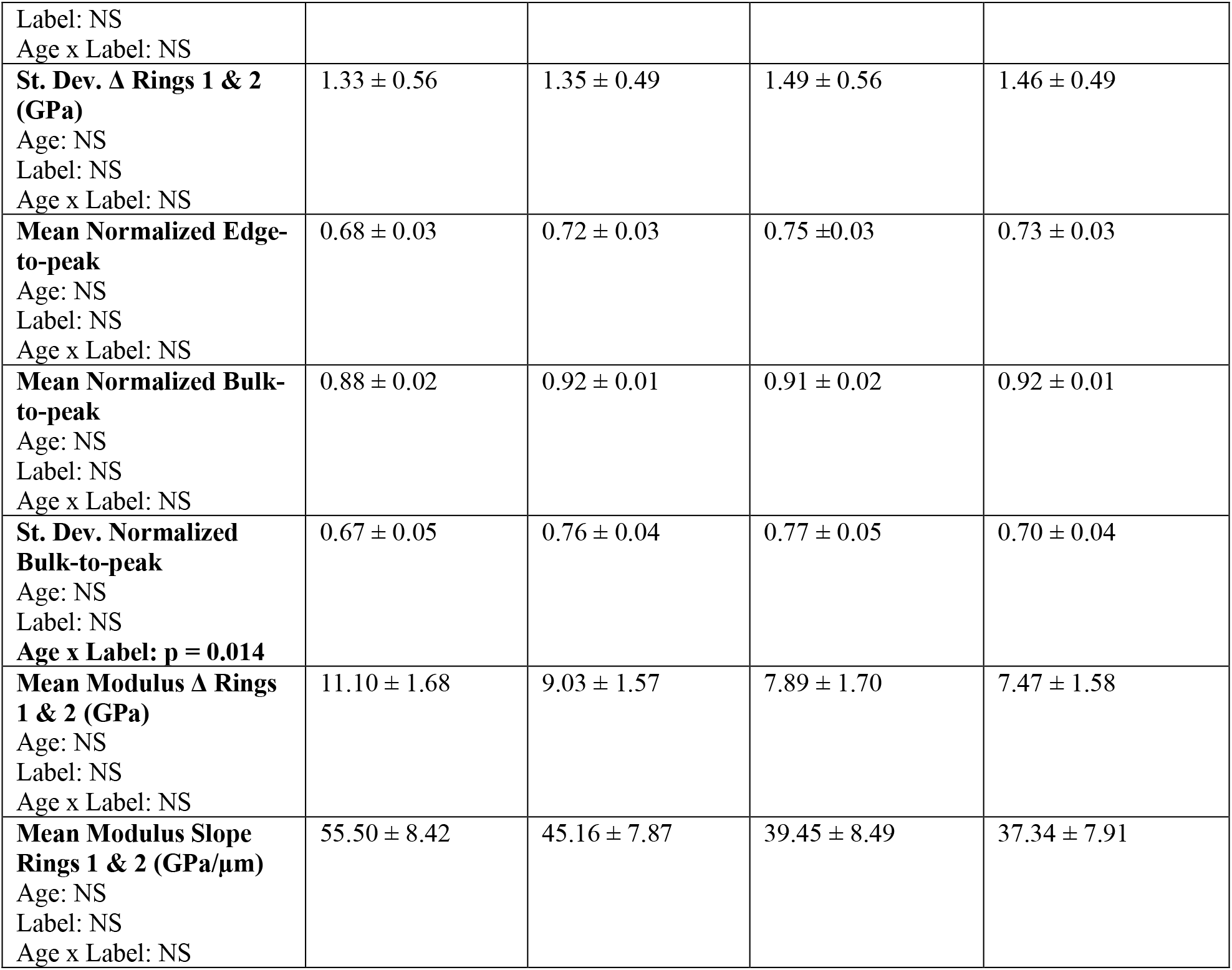
Measurements of lacunar morphological and modulus properties for 5 mo and 22 mo mice for bone-forming and non-bone-forming lacunae: Data are presented as marginal mean (adjusted for age and label) ± standard error. Bolded text indicates a statistically significant measure (p < 0.05). Values obtained through performing a mixed-model ANOVA.

**Figure 5.**
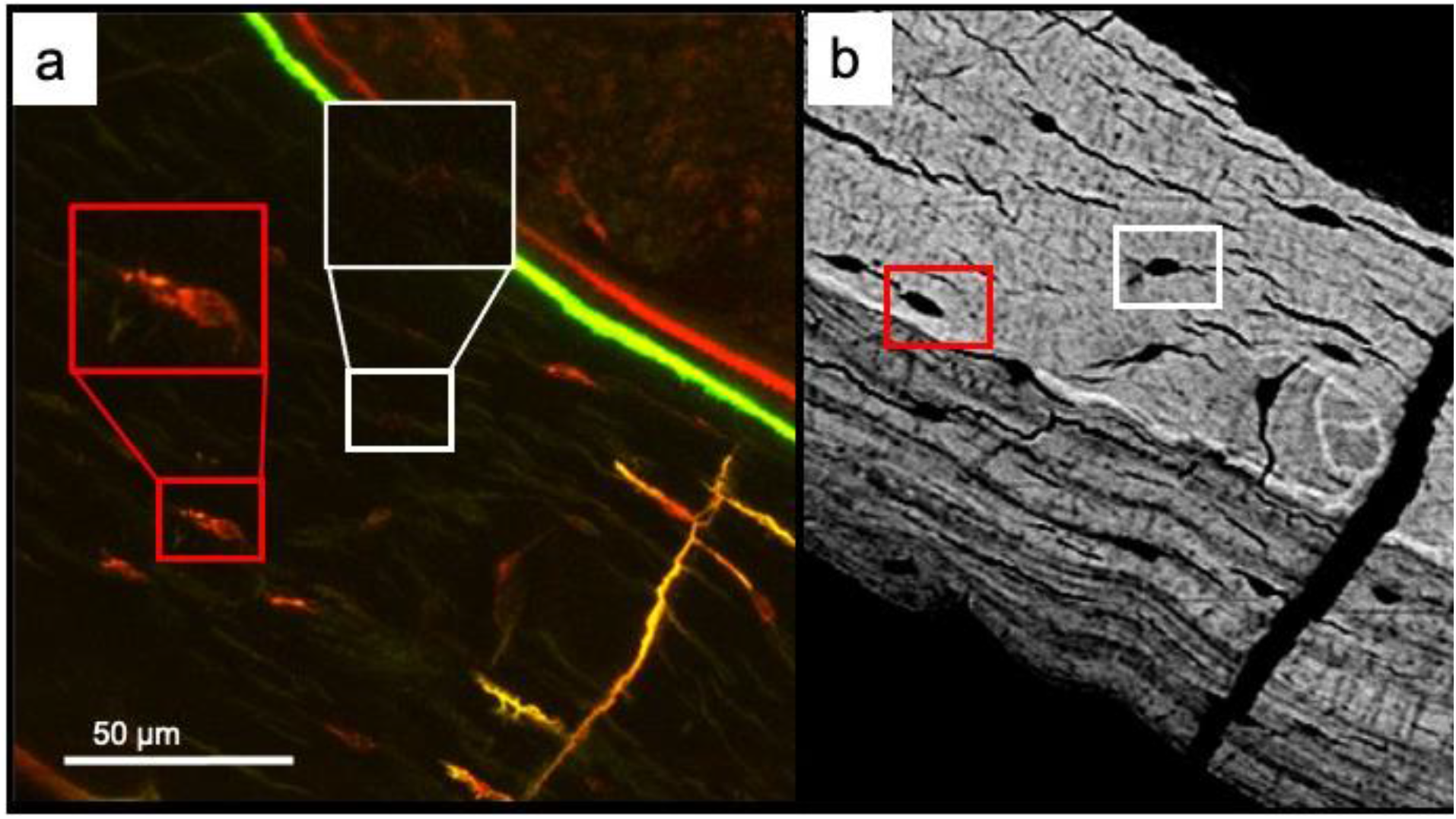
Remodeling osteocytes (red box: alizarin labeled lacuna) versus non-remodeling osteocytes (white box: non-labeled lacuna) imaged with a) confocal laser microscopy (63x-water immersion objective) and b) scanning electron microscopy (carbon coated surface, QBSD mode, 15 kV, 400x), both digitally zoomed.

Label did not significantly affect other measures of modulus gradation, including slope of the lacunar edge to peak modulus and the slope from the peak modulus to bulk modulus (**Table 1**). Measures of lacuna size, including area, minor and major axis length, and sphericity (minor / major axes) were not different between labeled and non-labeled lacunae. Additionally, there were no significant interactions with labels and aging for these measures.

### Early old age does not affect perilacunar modulus gradation but does affect peak-to-bulk modulus gradation and major axis length

The major axis of lacunae was smaller for 22 mo compared with 5 mo mice (−11.93%, p < 0.05), although area and minor axes where not changed (we note that major and minor axis lengths were determined through obtaining an elliptical fit for each lacunae, while area was determined through the number of pixels thresholded out by the MATLAB image processing code). Age did not impact most measures of modulus gradation. However, mean slope peak-to-bulk was significantly impacted by age; 22 mo mice showed a more gradual decrease in mean modulus from peak mean to bulk bone mean (−30.05%, p < 0.05, **Table 1**). There was a significant interaction between age and label for the bulk standard deviation normalized to the peak (p < 0.05). This measure evaluates the difference in heterogeneity of bulk bone compared to near the lacunar edge (typically where the maximum standard deviation occurs). This interaction is driven by increased normalized standard deviation for labeled compared with non-labeled lacunae for young mice. However, the p-value (+13.35%, p = 0.040) is not small enough to be considered a significant difference given our Bonferroni correction for family-wise error.

### Lacunar size and shape do not strongly correlate with perilacunar modulus gradation

Lacunar size and shape change in aging and are commonly studied with a variety of imaging techniques (e.g., CLSM, computed tomography techniques). Therefore, it would be useful to understand whether lacunar morphology can be used as an indication of perilacunar bone quality. We evaluated the strength of relationships between lacunar size and measures of modulus gradation (**Table S1**). The strongest Pearson correlations were major axis length vs. mean normalized edge-to-peak, minor axis length vs. normalized area before peak mean, and osteocyte area vs. normalized area before peak mean (r = −0.427, −0.341, −0.315, respectively). These results demonstrate that lacunar geometry is overall not a strong indicator of measures related to perilacunar modulus gradation.

## DISCUSSION

The osteocyte lacunar-canalicular system (LCS) is an important contributor to systemic mineral homeostasis. Osteocytes can remove and replace bone mineral around the expansive LCS surface (i.e., LCS remodeling). However, important questions remain about the impact of LCS remodeling on the quality of bone tissue surrounding this network. Aging truncates LCS morphologies [19,20,25,27,46] and increases the prevalence of osteocyte apoptosis and senescence [24,47–49]. Therefore, we were motivated to investigate whether osteocyte perilacunar bone quality differs between bone-forming and non-bone-forming osteocytes, and whether perilacunar bone quality differs in aging. We utilized confocal laser scanning microscopy and atomic force microscopy to evaluate gradations in perilacunar bone modulus around bone-forming and non-bone forming cortical femur osteocytes for skeletally mature young adult (5 mo) and early old age (22 mo) C57Bl/6 females.

Because synchrotron radiation studies show graded bone mineralization within hundreds of nanometers near lacunar and canalicular surfaces [1,4,33,50–52], we first determined the resolution at which we could resolve gradation in perilacunar moduli. We used AFM to map modulus for 12 μm x 12 μm areas surrounding lacunae (512 x 512 points, ~ 20 nm resolution) and developed an analysis procedure to assess mean modulus at distance increments from the lacunar wall. Using the same maps from an initial set of osteocyte scans from a young adult female C57Bl/6 mouse, we investigated modulus gradation from the lacunar wall at 0.2 - 1 μm step sizes outwards to 2 μm from lacunae (**Figure 3**). Our data indicate that at 0.2 μm resolution, an increase in modulus to a peak value is apparent, usually within 0.2 – 0.4 μm from the lacunar wall. At either 0.5 or 1 μm step size, these peak values are not resolved. However, decrease in bone tissue variability from the lacunar wall was resolved at all three step sizes. Importantly, many common bone quality assessment techniques (e.g., quantitative backscattered SEM, Raman spectroscopy, nanoindentation) do not have adequate resolution to observe mean perilacunar modulus gradation except for circumstances with large perturbations to mineral homeostasis [6,14,40,42,53–56].

Our results demonstrate that a substantial quantity of bone is likely impacted by osteocyte remodeling. The area of perilacunar bone with lower modulus, estimated from the average perilacunar bone area before the peak modulus for labeled lacunae, was 16 μm^2^. Because many (~60%) of our randomly-selected lacunae were forming bone, we can estimate that a sizable surface of bone along the expansive LCS has lower modulus associated with osteocyte bone formation. Importantly, perilacunar modulus gradations were only moderately correlated with lacunar geometry. Thus, morphological techniques are not sufficient for assessing changes to perilacunar bone quality.

The gradation of bone tissue modulus with distance from the lacunar wall was in excellent agreement with synchrotron studies studying gradation in mineral near the LCS in human [33] and ovine bone [57]. Hesse and co-authors studied lacunae and canaliculi from human mandible and found an immediate increase in mass density moving away from lacunar and canalicular edges to a peak at about a 0.2 μm distance from the lacunar edge. These peak values were followed by a decrease in mass density as measurements approached bulk bone tissue [33]. Most of our modulus measurements matched these profiles. In another study, Nango and colleagues assessed sub-micrometer bone mineralization gradients surrounding both osteocytes and canaliculi through use of a combination of synchrotron x-ray microscopy and transmission electron microscopy (TEM). The mineralization profiles observed for wild-type and osteoporotic mice showed gradations that shared an overall similarity with our study. The lowest mineralization was adjacent to the lacunar wall and increased with distance to either a peak or an asymptotic value [1]. Together, these findings suggest that our observed bone tissue modulus gradation is likely associated with variation in bone mineralization.

Perilacunar bone gradation may be influenced by a combination of active and passive mineralization and demineralization processes. We observed bone modulus gradations in 2D space around both labeled and unlabeled osteocytes, though the size and shape of these gradations depended on whether the osteocyte lacuna was fluorochome-labeled. Mineral exchange may be active since osteocytes are known to participate in mineral homeostasis [5,21,32,59] and in this process can acidify bone matrix and form osteoid [5,10,14,26,58]. Mineral exchange could also exist as a passive process as calcium exchange occurs between bone local to the LCS and interstitial fluid [33]. As suggested by Hesse and co-authors, mass density gradients followed by a peak may indicate a diffusion limit for calcium ions from LCS into the extracellular matrix (ECM) [33]. Alternatively, our observed modulus gradation and peak modulus values may also represent a spatial limit for lacunar bone tissue dissolution and re-mineralization by osteocytes. It is possible that the activities of passive as opposed to active mineralization changes with aging and in disease processes where osteocyte viability declines, but addressing this question requires further investigation.

A challenge in osteocyte research is relating the behavior of individual osteocytes with the impacts to the surrounding bone tissue material. This connection remains elusive, in part because the fixation and decalcification necessary to assess parameters of osteocyte behavior (e.g., apoptosis, senescence) generally precludes the determination of bone material properties. While Hesse and co-authors did not link bone mineral gradation with the health or activity of individual cells, they did observe altered mass density profiles in osteocytes from BRONJ mandibles [33]. For these specimens, in which osteocyte apoptosis is expected to be more common, peak mass density was higher and decay to bulk mass density was less, suggesting a mineral saturation effect. In the present work, we introduce a strategy to evaluate material properties around individually identified bone-forming osteocytes. We identified that fluorochrome bone labels administered 2d before euthanasia abundantly label osteocytes in the murine cortical femur at both 5 mo and 22 mo. The prevalence of alizarin and calcein labels administered 2d before euthanasia is similar, suggesting that the labeling is not specific to one label chemistry (**Figure S1**). Labels administered at 8d before euthanasia are infrequently observed, demonstrating that between 2d and 8d, labeled bone is often broken down by either active (i.e., osteocyte bone resorption) or passive (i.e., bone demineralization in contact with extracellular fluid) demineralization. These bone labels are easily observed using confocal laser scanning microscopy and the same lacunae can be mapped with atomic force microscopy. Thus, it is possible to test whether osteocytes forming bone have different perilacunar bone quality than osteocytes that are not forming bone.

Perilacunar modulus gradation was influenced by osteocyte bone formation but not by age. The lack of bone tissue modulus change seen in aging is consistent with other studies, where cortical bone nanoindentation modulus did not differ between young adult (4-6 mo) and early old age (19-24 mo) male C57Bl/6 mice [59,60]. We also evaluated whether perilacunar bone is less heterogeneous with increased age. Whereas perilacunar bone was always more variable near the lacunar wall, it was not differently variable either near or far the lacunar wall in aging. Thus, our evidence does not support that disruption to heterogeneity at this length scale is associated with age-related differences in bone quality [6,61]. Instead, other tissue toughening mechanisms, such as energy dissipation of less mature tissue, may be more important to the decline of bone fracture resistance in aging [25,46,62]. Overall, the specific impacts of perilacunar remodeling on bone tissue toughening mechanisms would benefit from further investigation.

Our study has several limitations. First, bone samples were dehydrated in ethanol and embedded in PMMA. Bone tissue dehydration and embedding each stiffen bone but do not disrupt bone mineralization [63]. In our MATLAB-based segmenting and thresholding procedure, we excluded measurements <5 GPa since these values would be indistinguishable from PMMA, and measurements >90 GPa as these values are most likely a result of alumina beads embedded in the sample during polishing. Mapping hydrated bone samples would reveal bone material properties closer to those *in vivo*. Another limitation is that pericanalicular bone tissue was not mapped in this study. In synchrotron studies, bone mass gradation around canaliculi is similar to around lacunae [33], suggesting that AFM may also resolve modulus gradation around these structures. The approach presented herein could be readily modified to map modulus around canaliculi or dendrites. We did not evaluate whether aging decreases the number or proportion of labeled lacunae. However, of all the randomly selected lacunae in this study, a similar proportion were labeled for young adult and early old age mice. Finally, this study assessed a small number (n = 5 / group) of young adult and early old age C57Bl/6 female mice. While we observed modulus gradation for every perilacunar bone map acquired for both ages, the causes of changes of modulus profiles with age would benefit from additional mice of both sexes across an extended age range.

We report, for the first time, that bone modulus is graded at the sub-micrometer scale around osteocyte lacunae. Perilacunar gradation is distinct for bone-forming lacunae for both young adult and early old age mice and in both cases is consistent with decreased tissue maturity. Given the immense scale of the LCS and abundance of osteocyte bone formation, our findings support the possibility that lacunar-canalicular remodeling can impact bone quality.

## METHODS

### Animal models

This investigation was conducted in two studies. The first study, in which methods were developed for AFM perilacunar bone modulus analysis, a 7-month-old female C57Bl/6 mouse was obtained from a live animal colony (group housed, 3 mice per cage, standard rodent chow and water provided *ad libitum*) at Montana State University. This mouse was euthanized via CO2 inhalation. The second study, which evaluated the effects of age and label on perilacunar modulus gradation, included 5-month (n = 5) and 22-month-old (n = 5) female C57Bl/6 mice from Charles River Laboratory. These mice were group housed (2-5 per cage), fed low fat diet (Research Diets D12450H; 10% kcal from fat) *ad libitum* for 8 weeks prior to euthanasia as controls for another study, provided water *ad libitum*, and euthanized via isoflurane inhalation. All animal procedures were approved by the Montana State University Institutional Animal Care and Use Committee.

### Sample preparation

Left femurs were harvested and fresh frozen at −20° C immediately after euthanasia. Femurs were gently thawed and tested to failure in three-point bending (results reported in a separate study). The distal halves of the femurs were histologically dehydrated in a graded ethanol series and embedded in poly(methyl) methacrylate (PMMA). Embedded distal femurs were sectioned at the midshaft using a low-speed diamond saw (Isomet, Buehler, Lake Bluff, IL), to obtain a transverse section with a 5 mm thickness. Then, cortical surfaces were polished with 600 and 1200 grits of wet silicon carbide papers (Buehler, Lake Bluff, IL), followed by fine polishing with Rayon fine clothes and different grades of alumina pastes (9, 5, 3, 1, 0.5, 0.3, and 0.05 μm) to achieve a mirror-like finish. Between polishing steps, sections were sonicated in tap water to remove any remaining particles. Embedded femur sections were mounted on a metal disk using epoxy (MasterBond EP29, Hackensack, NJ). A glass slide of the same 5 mm height was mounted next to the embedded femur section to be used for tip radius calibration.

### AFM mapping

Atomic force microscopy (AFM) analyses were performed with an Asylum Research Cypher S force microscopy system with an etched silicon tip (RTESPA-525, 200 N/m spring constant, Bruker AFM Probes, Camarillo, CA). AFM was operated in two different modes: AC tapping mode (for topography scans) and fast force mapping (for modulus maps). Using AC tapping mode, the cantilever was driven at a constant amplitude at its resonance frequency and scanned across the surface to measure topography of the bone samples and to locate lacunae. Fast force mapping generated an array of local force-distance curves, obtained at high speed with nanometer spatial resolution and was used to characterize modulus profiles around lacunae. Tip parameters were calibrated and resulting force curves were fit to a Hertzian contact model to calculate the contact modulus of the material [64]. First, calibration of a cantilever spring constant was obtained *via* thermal tune. Next, a force-distance curve was performed on a silicon wafer (Silicon inc., Boise, ID) to calculate optical lever sensitivity. Once these values were obtained, tip radius was calibrated by first acquiring a fast force map (320 x 320-pixel map) of a glass surface with known modulus (72 GPa, Fisherbrand, Pittsburgh, PA) then identifying the tip radius value needed to generate agreement of the Hertz model with the glass calibration surface.

For each bone, lacunae were randomly selected from the anterior side of the midshaft cortical cross-section. Selected lacunae were at least >20 μm away from bone endocortical and periosteal surfaces. A topographical lacunar map was first generated using AC mode, then fast force mapping generated a 512 x 512-pixel (12 x 12 μm map size at scan rate of 300 Hz) of lacuna with a ~ 20 nm resolution. A threshold of 500 nN was found to provide sufficient signal to noise ratio in the force curves and good agreement with the Hertzian contact models. While force curves represent an intermittent contact, rather than continuous contact technique, measurements of modulus must still account for potential tip wear. Tip radius was calibrated both before and after every fast force map of bone tissue was obtained, and the mean value of tip radius input into the Hertz model for modulus calculations. Tip radii were kept between 10 nm (pristine) and 20 nm, as these values are consistent with typical tip wear in literature and are smaller than the resolution of acquired modulus maps [65,66]. For reliability, we considered larger values to be the result of either breaking or contamination of the probe tip and associated data was not considered.

### Importing data and identifying the lacunar edge

Initially, square maps (equal number of x and y pixels) are imported to MATLAB as .csv files. The lacunar edge is defined for each map and points within the lacuna are masked out. Because dendrites extend from the lacunar wall, erosion is necessary to define a close-fitting lacunar edge. Erosion is performed based on a diamond-shaped element with size specified by the user (i.e., larger elements yield more aggressive erosion). The results of this step are shown in **Figure S2**. This step creates an array of points that describes the lacunar boundary. This process is repeated to create an over-eroded boundary. This over-eroded boundary will be utilized later in the code to create sequential boundaries. An over-eroded boundary is required due to an inherent dilation when using a smoothing function later in the code.

### Map and edge rotation

Next, maps are rotated about the lacunar centroid so that the ellipsoidal long axis of the lacuna is vertical. This step reduces distortion of sequential boundaries during dilation steps later in the code.

### Creation of sequential boundaries

The lacunar edge boundary created from the over-erosion step will be used to create sequential boundaries (e.g., separated by a specified distance) surrounding the lacuna. User inputs include the number of desired dilations (e.g., number of sequential regions of interest with increasing distance from the lacunar wall), the distance between each boundary, and the map dimensions. The results of this step are ‘unsmoothed’ boundaries, as shown in **Figure 1b**.

### Smoothing of sequential boundaries

The sequential boundaries are then smoothed via a convolution matrix [67]. This achieves a boundary that closely matches lacunar geometry but removes more harsh edges and features that need not be considered. However, this step inherently dilates the lacunar edge somewhat, hence over-erosion is necessary (***Identifying the lacunar edge***) in pre-processing. The results of the smoothing are shown in **Figure 1b** and **1c**.

### Binning and analyzing points between concentric boundaries

Next, points within each two sequential boundaries are binned. The x-y position of each pixel is matched with a corresponding modulus. Lastly, a histogram is created for each region using a discrete range of 1 GPa for histogram bin sizes (for example, if the range of the points within a certain region is 5.7 to 34.2 GPa there would be a bin for 5-6, 6-7, etc. up to 34-35). Several statistical measurements are made for each concentric regions including, range, median, mean, standard deviation, and full width at half maximum.

### Analysis of modulus versus distance from the lacunar wall

Using measures calculated from histograms, modulus versus distance profiles were generated. From these profiles, measures included the peak modulus (greatest mean modulus of all concentric regions versus distance from the lacunar wall), the bulk modulus (the mean modulus of the last concentric ring, 1.8-2 μm), the difference between the peak and bulk measures, the edge-to-peak and peak-to-bulk slopes (GPa/μm), and perilacunar area before peak modulus. Additionally, the slope measures were also calculated after normalizing to the peak modulus of a given map (**Figures 1d–1f**).

### Confocal laser scanning microscopy imaging

Samples were imaged using an upright confocal microscope (Leica SP3,) with the following parameters: 40x immersion lens, laser wavelength excitation of 488 nm (emission length 502-540) for calcein label and 561 and 633 nm (emission length 580-645) for alizarin label, pinhole set at 1 Airy unit, 1024 × 1024 resolution with a 600 Hz speed, and laser intensity set at 30% of the full power. The gain and offset were chosen such that in the images acquired the lacunae and their perilacunar remodeling were visible with minimum amount of noise.

### Statistical methods

Mixed-model ANOVA evaluated the impact of the fixed effects of label (label vs no label) and age (5 vs 22 mo) and the random effect of individual mouse on measures pertaining to modulus variation near lacunae (e.g., peak modulus, bulk modulus, etc). Residuals for all models were checked for normality and equal variance. The dependent variable was natural log transformed, if necessary, to satisfy these assumptions. Significance was defined *a priori* as p < 0.05. In the case of a significant interaction between age and label, post-hoc tests were adjusted for family-wise error with the Bonferroni procedure (i.e., 2 comparisons: label vs non-labeled at each age; critical α adjusted to p < 0.025). All analyses were performed using Minitab v.19.

## ASSOCIATED CONTENT

### Supporting Information

**Table S1.** Pearson correlations between lacunar size and shape measures vs. measures of modulus gradation.

| Correlations | Osteocyte Area<br>($\mu\text{m}$) | Major Axis<br>($\mu\text{m}$) | Minor Axis<br>($\mu\text{m}$) | Sphericity |
| --- | --- | --- | --- | --- |
| Location Peak Mean ($\mu\text{m}$) | -0.100 | -0.078 | -0.151 | -0.086 |
| $\Delta$ Mean (peak - bulk) | -0.178 | -0.054 | -0.105 | -0.039 |
| Mean slope Edge-to-peak (GPa/$\mu\text{m}$) | 0.185 | 0.301 | -0.026 | -0.229 |
| Mean Slope Peak-to-bulk (GPa/$\mu\text{m}$) | 0.125 | -0.006 | 0.094 | 0.074 |
| Mean Normalized Edge-to-peak | -0.304 | -0.427 | 0.060 | 0.383 |
| Mean Normalized Bulk-to-peak | 0.203 | 0.18 | 0.004 | -0.15 |
| Mean Modulus $\Delta$ Rings 1&2 (GPa) | 0.125 | 0.249 | -0.154 | -0.339 |
| Mean Modulus Slope Rings 1&2 (GPa/$\mu\text{m}$) | 0.125 | 0.249 | -0.154 | -0.339 |
| Area Before Peak Mean ($\mu\text{m}^2$) | 0.02 | 0.059 | -0.094 | -0.131 |
| Normalized Area before Peak Mean | -0.315 | -0.231 | -0.341 | -0.144 |

## AUTHOR INFORMATION

## Corresponding Author

Department of Mechanical & Industrial Engineering, Montana State University, P.O. Box 173800 Bozeman, MT 59717, United States of America Email address

## Author contributions

Experimental design: C.J.R., L.M.C., C.M.H. Data collection: C.J.R., G.V., A.D. Data analysis and interpretation: C.J.R., C.M.H. Manuscript drafting: C.J.R., C.M.H. Approval of final manuscript: all authors.

## Acknowledgements

This work was supported by NIH R03 AG068680, NSF 2120239, and NIH P20 GM103474. This work was performed in part at the Montana Nanotechnology Facility, a member of the National Nanotechnology Coordinated Infrastructure (NNCI), which is supported by the National Science Foundation (Grant# ECCS-2025391). We thank Dr. Heidi Smith at the Montana State University Center for Biofilm Engineering for assistance with CLSM imaging.

## Abbreviations

LCS: lacunar-canalicular system
AFM: atomic force microscopy
CLSM: confocal laser scanning microscopy

## Data availability

MATLAB code data have been deposited in the GitHub repository: https://github.com/cjrux/Osteocyte_Boundary_Analysis

---

**Figure S1.**
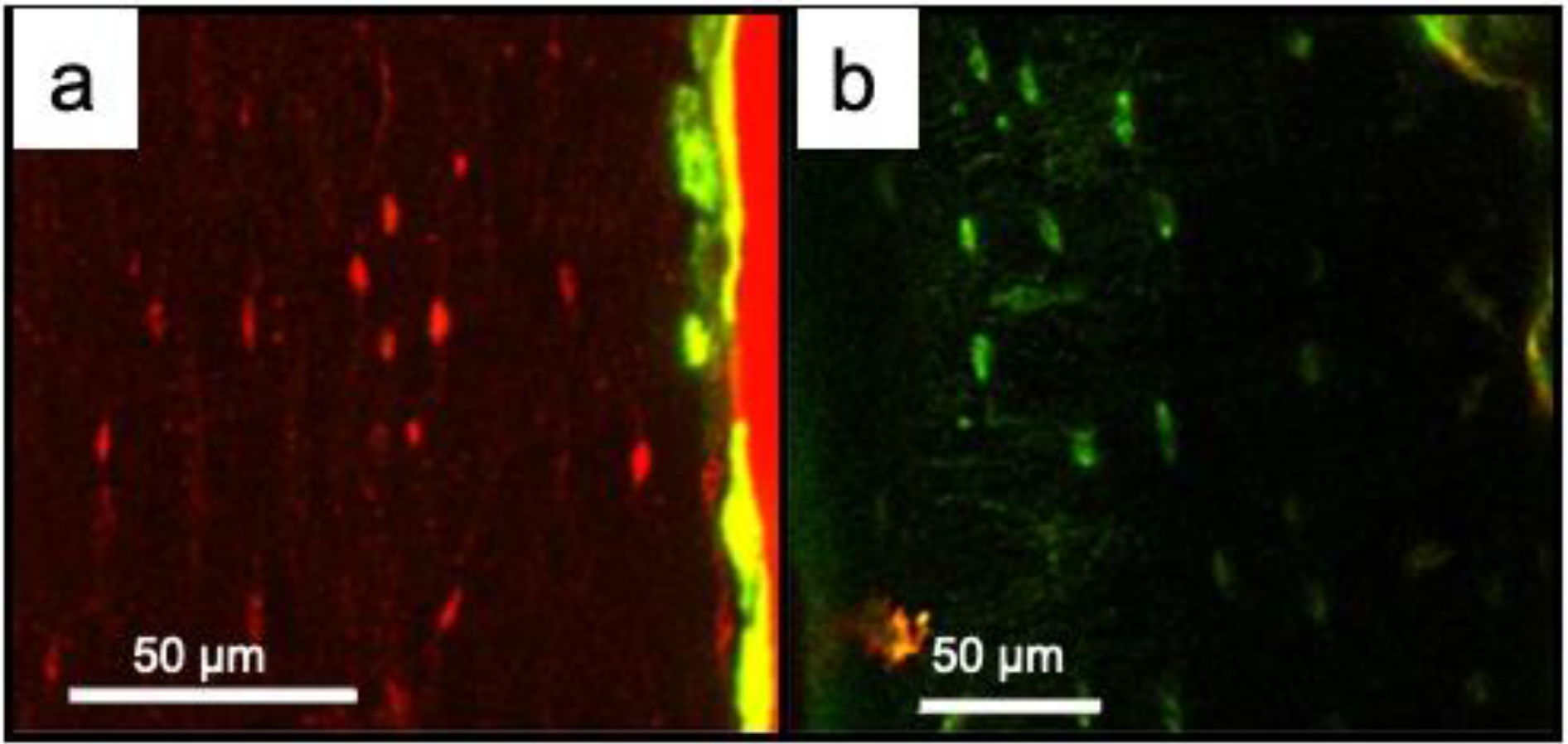
Osteocyte remodeling abundantly occurs shortly before euthanasia (2 days) regardless of the fluorochrome labeling order. a) Alizarin was administrated 2 days prior to euthanasia (calcein injection 6 days prior). b) Calcein was administrated 2 days prior to euthanasia (alizarin injection 6 days prior)

**Figure S2.**
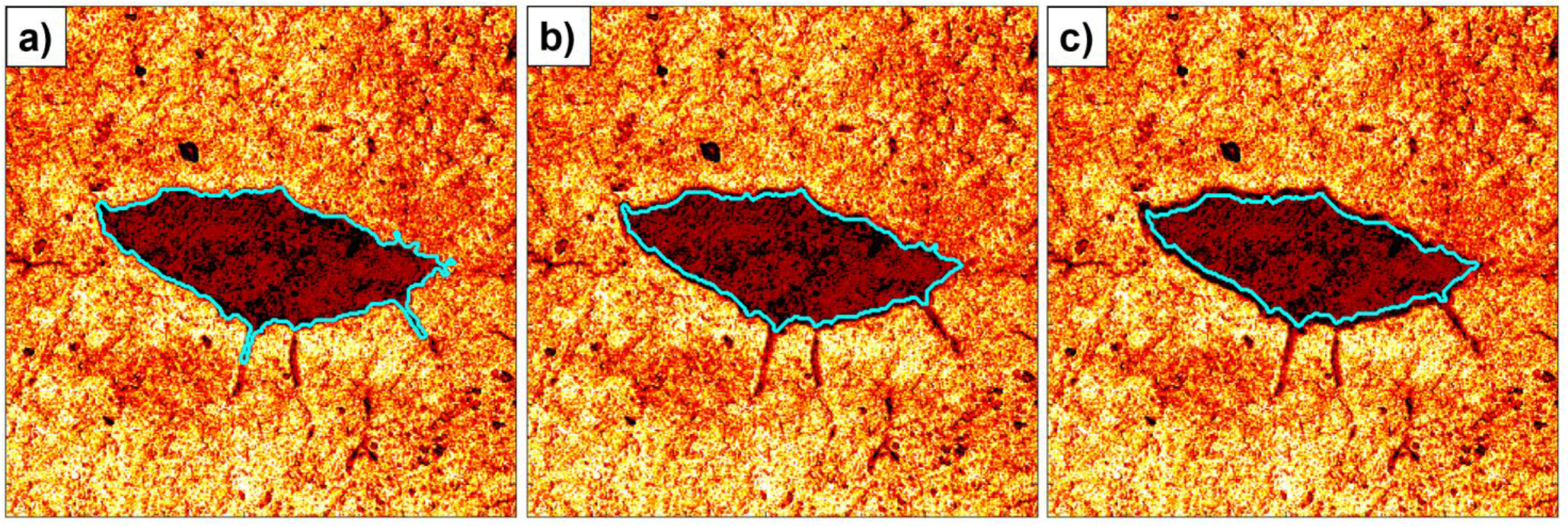
a) Under-eroded lacuna. b) Properly eroded lacuna. c) Over-eroded lacuna.

